# aPC/PAR1 confers endothelial anti-apoptotic activity via a discrete β-arrestin-2 mediated SphK1-S1PR1-Akt signaling axis

**DOI:** 10.1101/2021.03.27.437291

**Authors:** Olivia Molinar-Inglis, Cierra A. Birch, Dequina Nicholas, Metztli Cisneros-Aguirre, Anand Patwardhan, Buxin Chen, Neil J. Grimsey, Patrick K. Gomez Menzies, Huilan Lin, Luisa J. Coronel, Mark A. Lawson, Hemal. H. Patel, JoAnn Trejo

**Author notes:** For correspondence (JT).

## Abstract

Endothelial dysfunction is associated with multiple vascular diseases and lacks effective treatments. Activated Protein C (aPC) is a promising biotherapeutic that signals via protease-activated receptor-1 (PAR1) to promote diverse cytoprotective responses, including endothelial barrier stabilization, anti-inflammatory and anti-apoptotic activities, which is facilitated by co-receptors. We showed that aPC-activated PAR1 signals preferentially via β-arrestin-2 (β-arr2) and dishevelled-2 (Dvl2) scaffolds rather than G proteins to enhance barrier protection. However, the mechanisms by which aPC/PAR1 promotes other cytoprotective responses are poorly understood. Here we define a novel β-arr2-mediated sphingosine kinase-1 (SphK1)-sphingosine-1-phosphate receptor-1 (S1PR1)-Akt signaling axis that confers aPC/PAR1-mediated protection against cell death. We show that PAR1 and S1PR1 co-exist in caveolin-1-rich microdomains basally and aPC markedly increases S1PR1-caveolin-1 co-association. Moreover, aPC stimulates β-arr2-dependent SphK1 activation independent of Dvl2, which is critical for S1PR1 transactivation. These studies reveal that different aPC/PAR1 cytoprotective responses are mediated by discrete β-arr2-driven signaling pathways in caveolae.

## Introduction

Endothelial dysfunction, a hallmark of inflammation, is associated with the pathogenesis of vascular diseases and causes endothelial barrier disruption and increases sensitivity to apoptosis (Rajendran et al., 2013). There are limited treatment options for improving endothelial dysfunction, which is prevalent in diseases such as sepsis, a condition with high morbidity and mortality (Evans, 2018). Activated protein C (aPC) is a promising biotherapeutic that exhibits multiple pharmacological benefits in preclinical studies, including sepsis (Bernard et al., 2001; Griffin et al., 2015). In endothelial cells, protease-activated receptor-1 (PAR1), a G protein-coupled receptor (GPCR), is the central mediator of aPC cytoprotective responses, including endothelial barrier stabilization, anti-inflammatory, and anti-apoptotic responses (Bouwens et al., 2013; Mosnier et al., 2006; Shahzad et al., 2019). The pathways by which aPC/PAR1 elicits diverse cytoprotective responses are poorly defined.

APC-dependent endothelial cytoprotection requires the localization of PAR1 and the aPC co-receptor, endothelial protein C receptor (EPCR), in caveolin-1-rich microdomains (Razani et al., 2001). APC proteolytically activates PAR1 by preferential cleavage of the receptor’s N-terminal arginine (R)-46 residue, which is distinct from the canonical thrombin cleavage site at (R)-41 (Bae et al., 2007, 2008; Mosnier et al., 2012; Russo et al., 2009). APC/PAR1 signals preferentially via β-arrestin-2 (β-arr2), a multifunctional adaptor protein, and not heterotrimeric G proteins to promote cytoprotection (Kanki et al., 2019; Roy et al., 2016; Soh et al., 2011). We showed that aPC-activated PAR1 signals via β-arr2 and Dishevelled-2 (Dvl2) scaffolds to induce Rac1 activation and endothelial barrier protection (Soh & Trejo, 2011). β-arr2 and Dvl2 are also essential for aPC-mediated inhibition of cytokine-induced immune cell recruitment, an anti-inflammatory response (Roy et al., 2016). APC/PAR1 also stimulates Akt signaling and protects against endothelial cell death induced by tumor necrosis factor-alpha (TNF-α) and staurosporine; however, the role of β-arr2 and Dvl2 scaffolds in eliciting these specific anti-apoptotic responses is not known (De Ceunynck et al., 2018; Mosnier et al., 2003).

The interaction of GPCRs with co-receptors can alter the active conformation of receptors, β-arrestin recruitment, and biased signaling (Lee et al., 2020; Shen et al., 2018) and is relevant to aPC/PAR1-driven endothelial cytoprotective signaling. APC-activated PAR1 cooperates with PAR3 and sphingosine-1-phosphate receptor-1 (S1PR1) to drive cytoprotective signaling (Burnier et al., 2013; Feistritzer et al., 2005; Finigan et al., 2005). APC preferentially also cleaves PAR3 within the N-terminus at a non-canonical (R)-41 site to promote endothelial barrier protection *in vitro* and *in vivo* (Burnier & Mosnier, 2013). In contrast to PAR3, aPC signals indirectly to S1PR1 to enhance basal endothelial barrier stabilization and to protect against barrier disruption (Finigan et al., 2005)(Feistritzer & Riewald, 2005). However, the mechanism by which aPC/PAR1 transactivates S1PR1 and the role of S1PR1 in other aPC-mediated cytoprotective responses, such as anti-apoptosis, is not known.

In this study, we assessed whether S1PR1 and the β-arr2 and Dvl2 scaffolds function as universal mediators of aPC/PAR1 cytoprotection by examining anti-apoptotic responses. Using a combined pharmacological inhibitor and siRNA knockdown approach in human cultured endothelial cells, we define a novel β-arr2-sphingosine kinase-1 (SphK1)-S1PR1-Akt signaling axis that is necessary to confer aPC/PAR1-mediated protection against TNF-α-induced cell death. Our studies further demonstrate that aPC-stimulated phosphorylation, translocation, and activation of SphK1 are dependent on β-arr2 and not Dvl2. In addition, we show that S1PR1 and PAR1 co-localize and co-exist in Caveolin-1 (Cav-1)-rich microdomains, and aPC-stimulated S1PR1-Cav-1 co-association is dependent on PAR1. These studies reveal that different aPC/PAR1 cytoprotective responses are mediated by discrete β-arrestin2-driven signaling pathways modulated by co-receptors localized in caveolae.

## Results

### PAR1 and S1PR1 colocalize in caveolae

Enrichment of GPCRs in caveolae augments cell signaling efficiency and specificity (Ostrom et al., 2004). Cav-1 is a structural protein essential for caveolae formation and modulates the activity of signaling molecules (Fridolfsson et al., 2014; Machleidt et al., 2000). Previous studies showed that PAR1 and EPCR localize to caveolae, which is required for aPC-stimulated Rac1 activation and endothelial barrier protection (Bae et al., 2007, 2008; Russo et al., 2009). This prompted us to investigate whether S1PR1 localized to caveolae in endothelial cells. The localization of endogenous S1PR1 and PAR1 in Cav-1 enriched fractions was examined in HUVEC-derived EA.hy926 cells using sucrose gradient fractionation. PAR1 segregated into Cav-1-enriched fractions as reported previously (Fig. 1A) (Soh & Trejo, 2011). S1PR1 was also detected in Cav-1-enriched fractions (Fig. 1A), suggesting that both PAR1 and S1PR1 reside primarily in caveolae in human cultured endothelial cells. Next, we examined co-association of S1PR1 with Cav-1 by immunoprecipitation. Endothelial EA.hy926 cells were stimulated with or without aPC, S1PR1 was immunoprecipitated, and Cav-1 was detected by immunoblotting. aPC induced a significant increase in Cav-1 interaction with S1PR1, compared to unstimulated cells or IgG controls (Fig. 1B, lanes 1-3). Interestingly, in PAR1-deficient EA.hy926 cells, aPC failed to induce S1PR1-Cav-1 interaction (Fig. 1B, lanes 4-6). These data suggest that PAR1 and S1PR1 are both localized in caveolae (Fig. 1C) and that PAR1 is required for aPC-induced S1PR1 co-association with Cav-1.

We next examined PAR1 and S1PR1 co-localization in endothelial cells using immunofluorescence confocal microscopy. Confocal microscopy imaging showed that both PAR1 and S1PR1 are localized at the plasma membrane in unstimulated endothelial cells and remain at the cell surface after aPC treatment, suggesting that neither PAR1 nor S1PR1 internalize after aPC stimulation (Fig. 1D). This observation is consistent with previous studies that showed aPC stimulation failed to promote PAR1 internalization in endothelial cells (Russo et al., 2009; Schuepbach et al., 2008). In contrast, thrombin induced internalization of PAR1 observed by the loss of PAR1 at the cell surface and loss of PAR1-S1PR1 co-localization at the plasma membrane. (Fig. 1D). Thus, PAR1 and S1PR1 co-exist in caveolae and remain at the cell surface after prolonged aPC stimulation (Fig. 1D). While S1PR1 has been implicated in aPC-mediated endothelial barrier stabilization (Feistritzer & Riewald, 2005; Finigan et al., 2005), the role of S1PR1 in aPC/PAR1 driven anti-apoptotic responses in endothelial cells is not known and was examined next.

**Figure 1.**
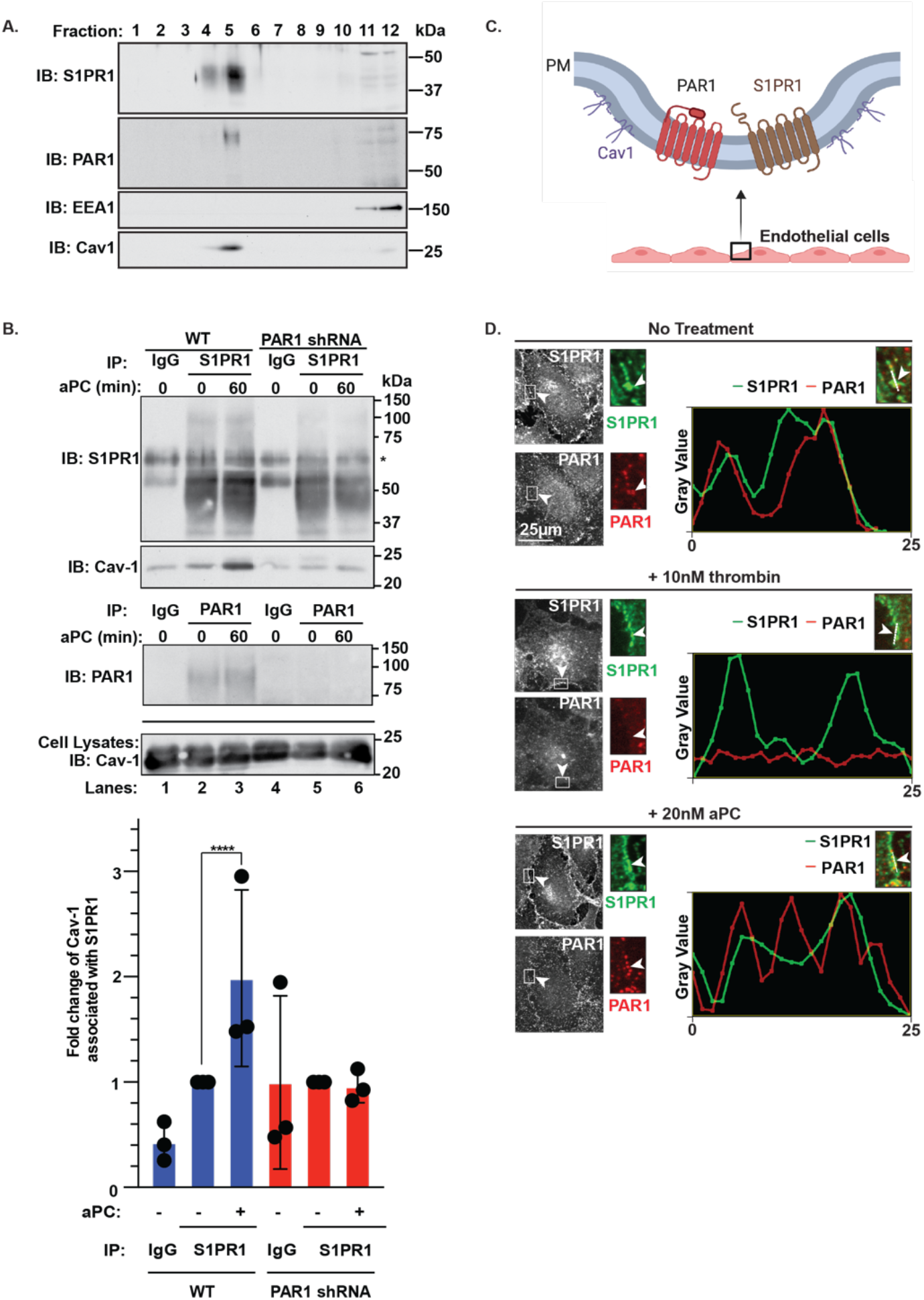
PAR1 and S1PR1 coexist in caveolae and colocalize at the plasma membrane. (A) EA.hy926 cells were lysed and subjected to sucrose gradient fractionation. Fractions were collected and immunoblotted for S1PR1, PAR1, EEA1, and Cav-1. (B) Wildtype (WT) and PAR1 shRNA expressing EA.hy926 cells were treated with or without 20nM aPC, lysed, immunoprecipitated and immunoblotted as shown. Data (mean ± S.D., n = 3) were analyzed by Student’s t-test (****, *P*<0.0001). (C) Schematic of endothelial cells expressing PAR1 and S1PR and localization in caveolae. (D) EA.hy926 cells untreated or treated with 10 nM ±-thrombin or 20 nM aPC for 60 min and immunostained for endogenous PAR1 (red) and S1PR1 (green) co-localization assessed by immunofluorescence confocal microscopy. Inset shows magnification of the cell periphery (white arrowhead) and white line drawn to show area analyzed for PAR1 and S1PR1 overlap. Line scan analysis was performed in ImageJ to assess PAR1 and S1PR1 overlap and plotted. Scale bar = 25 µm.

### S1PR1 is required for aPC-PAR1-mediated anti-apoptotic activity

To assess the role of S1PR1 in aPC/PAR1-driven anti-apoptotic responses, aPC-mediated protection against TNF-α-induced apoptosis was examined. Endothelial cells incubated with TNF-α for 20 to 24 h exhibited a significant increase in cell death as detected by Annexin V-FITC staining and flow cytometry, compared to untreated cells (Fig. 2A and B). In contrast, pretreatment with aPC for 4 h resulted in a significant reduction in TNF-α induced cell death, whereas incubation with aPC alone had no effect (Fig. 2A and B). Phase-contrast images of TNF-α treated endothelial cells display morphological signs of cell death, based on substantial cell shrinkage and rounding (Fig. 2A, *lower panels*), which is not detected in control cells and was reduced in aPC treated cells. Next, we determined if aPC treatment was sufficient to reverse TNF-α-initiated cell death by treating endothelial cells with TNF-α before aPC exposure. Similar to aPC pretreatment, post-treatment with aPC for 2 h or 3 h caused a significant reduction in TNF-α-induced cell death (Fig. 2C). To further assess the effect of aPC treatment on the TNF-α induced apoptosis, TNF-α-induced cleavage of Caspase-3, a key effector of apoptosis, was examined. TNF-α induced a significant increase in Caspase-3 cleavage (Fig. 2D, lanes 1-3), which was reversed by pre-and post-treatment with aPC (Fig. 2D, lanes 3-5). These results indicate that aPC confers protection against TNF-α-induced apoptosis.

**Figure 2.**
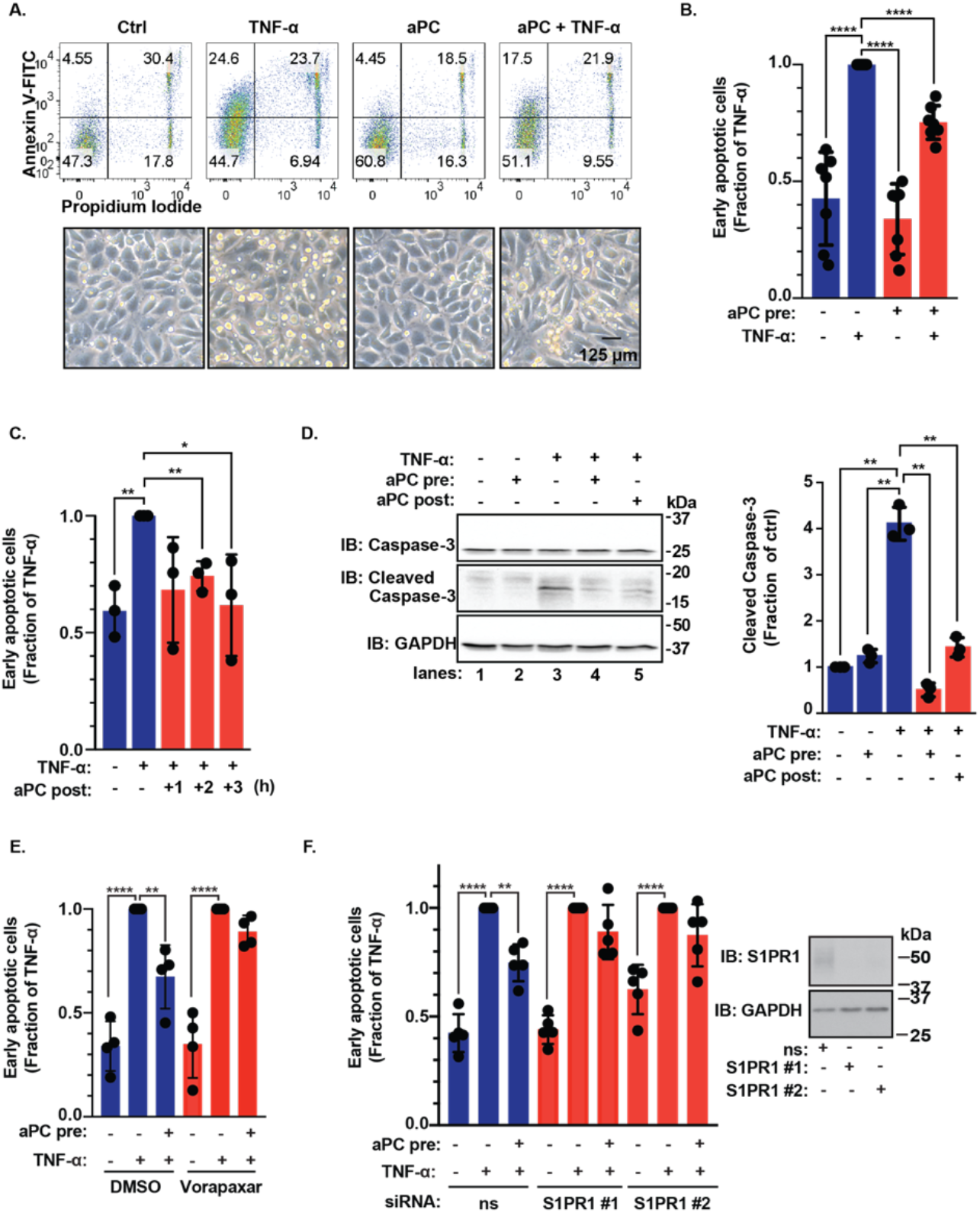
PAR1 and S1PR1 mediate aPC anti-apoptotic activity in endothelial cells. (A) EA.hy926 cells were pretreated with or without 20 nM aPC for 4 h and then treated with 20 ng/mL TNF-α for 24 h. A representative example of cell death determined by Flow cytometry using Annexin V-FITC and Propidium Iodide is shown in the top panel. Phase-contrast images of endothelial cells are shown before and after stimulation (*bottom panels*). Scale bar = 125 µM. (B) The data (mean ± S.D., n = 7) of the early apoptotic response (top right quadrant) were analyzed by Student’s t-test (****, *P*<0.0001). (C) EA.hy926 cells were treated with 20 ng/mL TNF-α for 24 h and then 20 nM aPC was added 1, 2, or 3 h after initial TNF-α stimulation. The data (mean ± S.D., n = 3) were analyzed by Student’s t-test (*, *P*<0.05; **, *P*<0.01,). (D) EA.hy926 cells were pretreated with or without 20 nM aPC for 3 h and followed by incubation with 25 ng/mL TNF-α for 24 h. Cleaved caspase-3 was detected by immunoblotting. Data (mean ± S.D., n = 3) was analyzed by Student’s t-test (**, *P*<0.01). (E) EA.hy926 cells were pretreated with 10 µM vorapaxar for 1 h. stimulated with 20 nM aPC for 4 h and then treated with 20 ng/mL TNF-α for 24 h and apoptosis determined. Data (mean ± S.D., n = 4) were analyzed by two-way ANOVA (****, *P*<0.0001; **, *P*<0.01). (F) EA.hy926 cells transfected with non-specific or two different S1PR1-specific siRNAs were stimulated with 20 nM aPC for 4 h, treated with 20 ng/mL TNF-α for 24 h. Cell lysates were immunoblotted to validate knockdown. Data (mean ± S.D., n = 5) were analyzed by two-way ANOVA (****, *P*<0.0001; \*\**P*<0.01).

Next, the role of PAR1 and S1PR1 in aPC-mediated protection against TNF-α induced apoptosis was assessed. PAR1 function in aPC-induced cell survival was examined using the PAR1 selective antagonist, vorapaxar, which effectively blocked aPC-mediated anti-apoptotic activities compared to control cells (Fig. 2E). SiRNA-targeted depletion of S1PR1 was utilized to assess the function of S1PR1 in aPC/PAR1-mediated anti-apoptotic responses. Depletion of S1PR1 was confirmed by immunoblot (Fig. 2F, *inset*). As expected, TNF-α-induced cell death was significantly reduced by aPC in non-specific siRNA control cells (Fig. 2F). In contrast, aPC failed to protect against TNF-α initiated cell death in S1PR1-depleted cells (Fig. 2F). Thus, S1PR1 is an important mediator of aPC/PAR1-induced anti-apoptotic activities in endothelial cells, however the mechanism by which S1PR1 facilitates aPC/PAR1-promoted cell survival is not known.

### Akt mediates aPC-induced anti-apoptotic activity via an S1PR1-dependent pathway

Akt is a critical regulator of cell survival in multiple cell types (Fujio et al., 1999; Los et al., 2009; Samakova et al., 2019). To determine the role of Akt in aPC-mediated anti-apoptotic responses, we used the Akt inhibitor MK-2206. In endothelial EA.hy926 cells, aPC stimulated an early increase in Akt S473 phosphorylation that was sustained for 90 min (Fig. 3A, lanes 1-6), similar to previous reports (Mosnier et al., 2012; Sinha et al., 2018). Pretreatment of endothelial cells with MK-2206 resulted in significant reduction of aPC-induced Akt S473 phosphorylation (Fig. 3A, lanes 7-12), confirming that MK-2206 effectively inhibits agonist-induced Akt activation. MK-2206 was then used to assess Akt function in aPC/PAR1-protection against TNF-α-induced cell death. Compared to control cells, aPC-mediated protection against TNF-α-induced cell death was significantly decreased in MK-2206-treated cells (Fig. 3B), indicating that Akt activity is required for aPC-induced pro-survival effects in endothelial cells.

**Figure 3.**
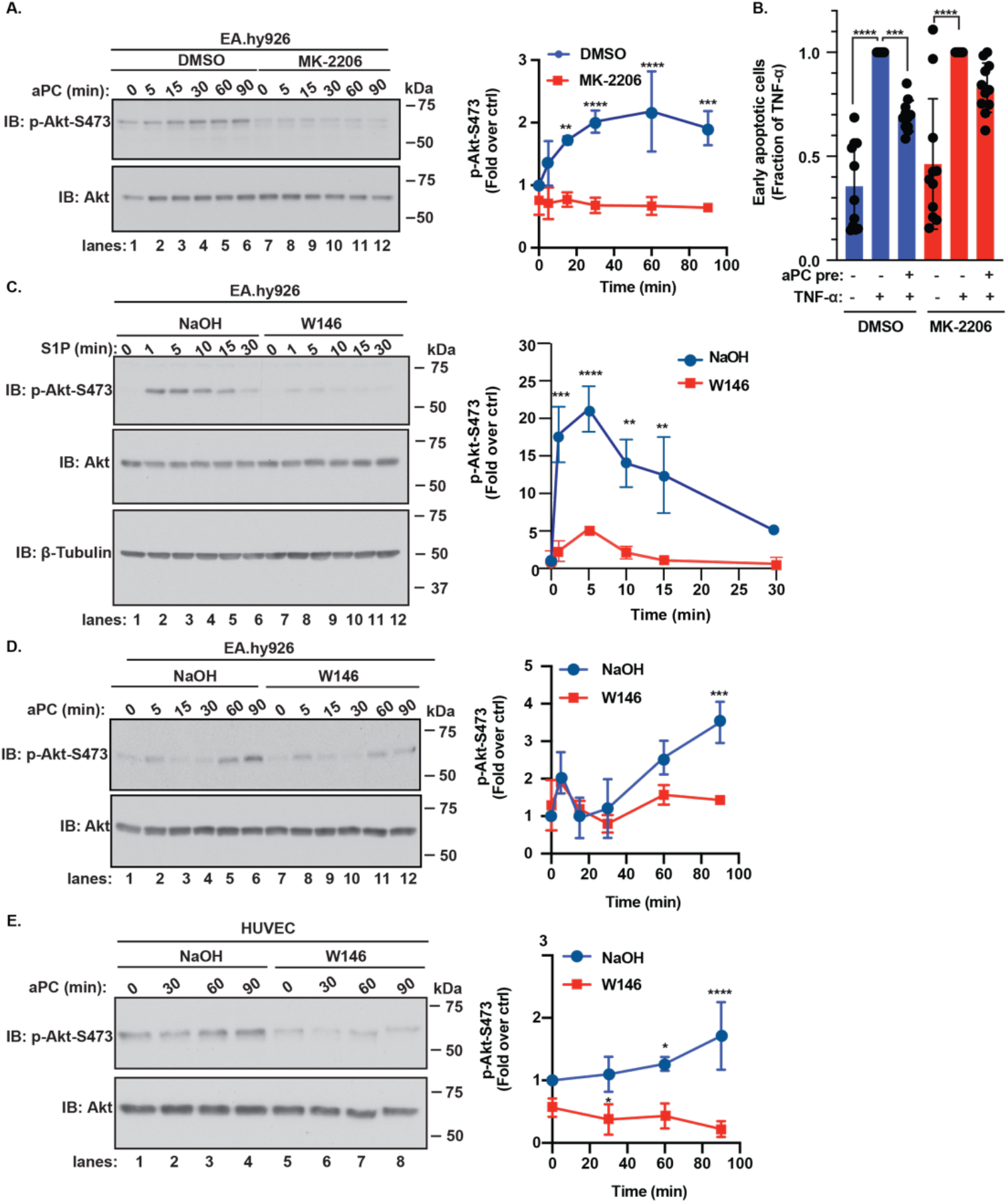
Akt mediates aPC anti-apoptotic activity through an S1PR1-dependent pathway. (A) EA.hy926 cells were pretreated with or without 1 µM MK-2206 for 1 h, stimulated with or without 20 nM aPC for various times, phospho-Akt-S473 was detected by immunoblotting. Data (mean ± S.D., n = 3) was analyzed by two-way ANOVA. (B) EA.hy926 cells were pretreated with or without 20 nM aPC for 4 h and then treated with 20 ng/mL TNF-α for 20-24 h and apoptosis measured. Data (mean ± S.D., n = 11) were analyzed by ordinary two-way ANOVA. (C) EA.hy926 cells were pretreated with or without 10 µM W146 for 30 min and with or without 1 µM S1P for a 30 min for various timesl phospho-Akt-S473 was then detected by immunoblotting. Data (mean ± S.D., n = 3) was analyzed by two-way ANOVA. (D) EA.hy926 cells were pretreated with or without 10 µM W146 for 30 min and with or without 20 nM aPC at different times, phospho-Akt-S473 was detected by immunoblotting. Data (mean ± S.D., n = 3) was analyzed by two-way ANOVA. (E) HUVECs were treated and data analyzed as described in D. ****, *P*<0.0001, ***, *P*<0.001, **, *P*<0.01, and \**P*<0.05.

The function of S1PR1 in aPC-induced Akt-dependent cell survival was next examined in endothelial cells using W146, a S1PR1 selective antagonist. Sphingosine-1-phosphate (S1P)-stimulated S1PR1 activation caused a rapid and significant increase in Akt S473 phosphorylation (Fig. 3C, lanes 1-6), which was significantly inhibited in W146 treated cells (Fig. 3C, lanes 7-12). In control-treated EA.hy926 cells, aPC caused a modest early peak in Akt phosphorylation followed by a more robust increase in Akt phosphorylation at later times (Fig. 3D, lanes 1-6). Inhibition of S1PR1 signaling with the W146 antagonist significantly reduced the late increase in aPC-PAR1-stimulated Akt activation but did not affect the early Akt response (Fig. 3D, lanes 7-12). A similar inhibitory effect of W146 on aPC-induced Akt phosphorylation detected at later times was observed in primary HUVECs (Fig. 3E). In contrast, inhibition of S1PR1 with W146 did not perturb aPC/PAR1-stimulated ERK1/2 activation in either EA.hy926 cells or HUVECs (Supplemental Fig. S1). These results demonstrate that selective antagonism of S1PR1 specifically blocks Akt activation and not ERK1/2 signaling. Thus, S1PR1 is required for aPC induction of Akt-driven cell survival signaling after prolonged agonist treatment. However, the mechanisms by which aPC/PAR1 transactivates S1PR1-Akt signaling have not been clearly defined.

### aPC-stimulated SphK1 activity is required for Akt activation

SphK1 catalyzes the phosphorylation of sphingosine to form S1P, the natural ligand that stimulates S1PR1 activation (Siow et al., 2011). SphK1 phosphorylation at serine (S)-225 is highly correlated with activation (Siow & Wattenberg, 2011) and was examined in aPC-treated endothelial cells using anti-phospho-SphK1-specific antibodies. APC induced a significant increase in SphK1 S225 phosphorylation at 5 min that was sustained for 120 min (Fig. 4A), consistent with prolonged aPC cytoprotective signaling (Feistritzer & Riewald, 2005; Finigan et al., 2005; Soh & Trejo, 2011). Activation of SphK1 through (S)-225 phosphorylation results in plasma membrane translocation (Johnson et al., 2002; Pitson et al., 2003; Pitson et al., 2005; ter Braak et al., 2009) and was examined using cellular fractionation. SphK1 was observed in both membrane and cytosolic fractions in unstimulated endothelial cells (Fig. 4B, lanes 1 and 3). However, aPC stimulated significant redistribution of SphK1 from the cytosol to the membrane fraction enriched in Na^+^/K^+^-ATPase, a plasma membrane resident protein (Fig. 4B, lanes 2 and 4). To determine if SphK1 S225 phosphorylation and translocation is associated with SphK1 activity, aPC-stimulated SphK1 activation was measured using a luminescence assay optimized to measure aPC-stimulated SphK1 activity (Supplemental Fig. S2A). APC induced a robust increase in SphK1 activity following 15 min of stimulation (Fig. 4C), which was significantly reduced by the SphK1 selective inhibitor PF-543 (Fig. 4C). Thus, aPC/PAR1 stimulates SphK1 S225 phosphorylation, translocation to the plasma membrane and activation.

**Figure 4.**
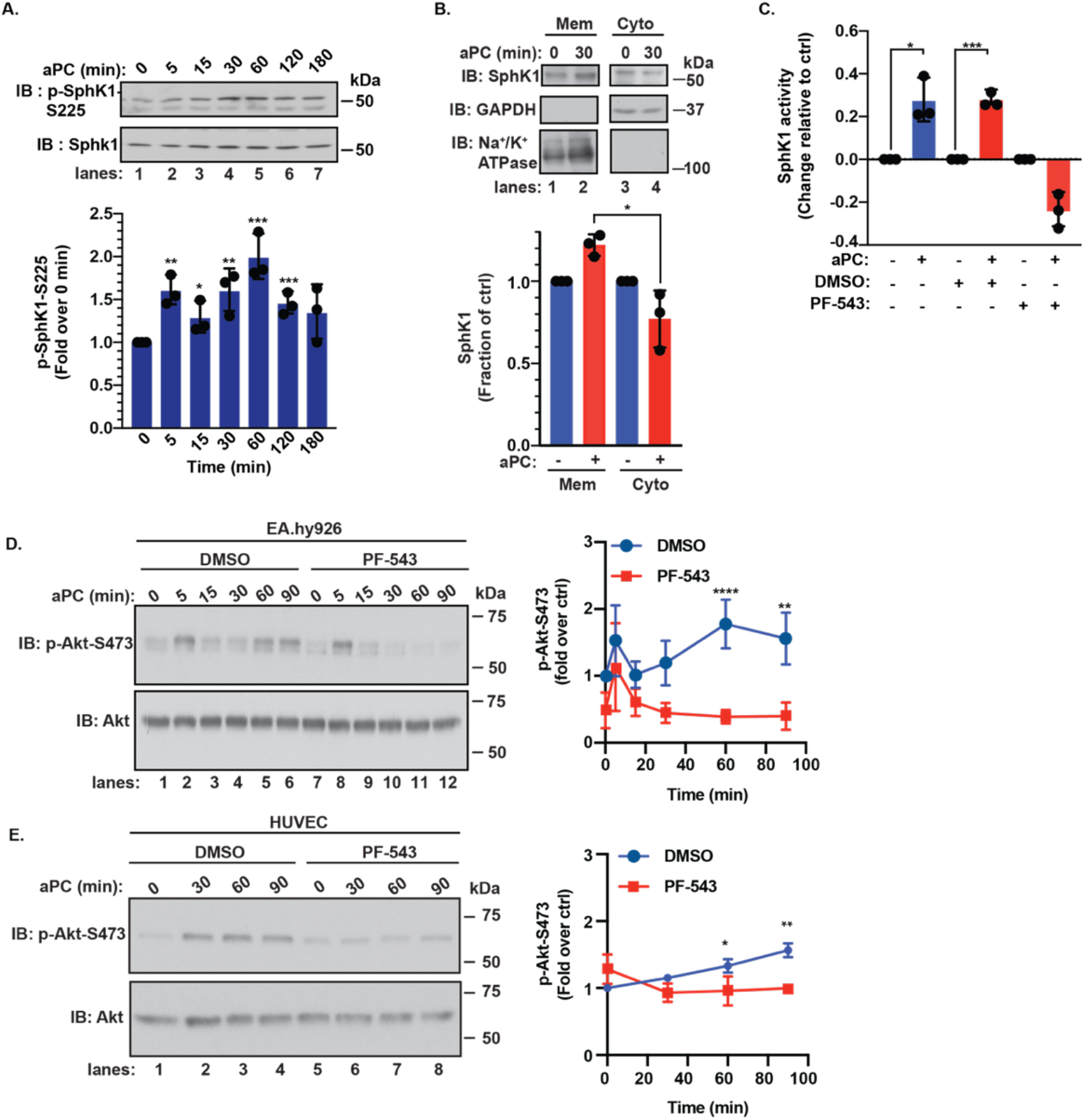
APC activates SphK1 and SphK1 is required for aPC-induced activity. (A) EA.hy926 cells were stimulate with or without aPC for 180 min, phospho-SphK1 was detected by immunoblotting. Data (mean ± S.D., n = 3) were analyzed by Student’s t-test. (B) EA.hy926 cells were stimulated with or without 20 nM aPC for 30 min, cells were lysed and subcellular fractionation performed. SphK1, the cytoplasmic control (GAPDH) and the membrane control (Na^+^/K^+^ ATPase) were detected by immunoblotting. Data (mean ± S.D., n = 3) was analyzed by Student’s t-test. (C) EA.hy926 cells were pretreated with or without 100 nM PF-543 for 1 h, and then treated with 20 nM of aPC for 15 min. Cells were lysed and SphK1 activity assessed by using the Echelon SphK1 activity assay. Data (mean ± S.D., n = 3) was analyzed by Student’s t-test. (D) EA.hy926 cells were pretreated with or without 100 nM PF-543 for 1 h and stimulated with or without 20 nM aPC for various times, phospho-Akt-S473 was detected by immunoblotting. Data (mean ± S.D., n = 3) was analyzed by two-way ANOVA. (E) HUVECs were treated and analyzed as described in D. ****, *P*<0.0001, ***, *P*<0.001, **, *P*<0.01, and *, *P* <0.05.

To determine whether SphK1 activity is linked to aPC/PAR1-S1PR1-dependent Akt activation, endothelial cells were treated with PF-543. In control endothelial EA.hy926 cells, aPC stimulated increased Akt phosphorylation at early and late times (Fig. 4D, lanes 1-6). However, aPC failed to induce Akt signaling at the late times in cells pretreated with PF-543 (Fig. 4D, lanes 11-12 versus 5-6), whereas the early increase in Akt phosphorylation was not perturbed (Fig. 4D, lane 2 versus 8). PF-543 similarly blocked aPC-induced Akt phosphorylation in HUVECs at late times (Fig. 4 E, lanes 1-4 versus 5-8). In contrast to Akt, aPC-induced ERK1/2 activation was not perturbed by PF-543 in either EA.hy926 or HUVECs compared to DMSO control cells (Supplemental Fig. S2B and S2C). These results suggest that aPC-stimulated SphK1 activity is specifically linked to transactivation of the S1PR1-Akt signaling axis.

### β-arr2 initiates aPC-induced SphK1-dependent S1PR1-Akt pro-survival signaling

β-arr2 and Dvl2 function as scaffolds and facilitate aPC/PAR1-induced endothelial barrier protection (Roy et al., 2016; Soh & Trejo, 2011). However, it is not known if β-arr2 and Dvl2 are similarly required for aPC-induced SphK1 activation and were examined using siRNA targeted depletion. SiRNA-mediated depletion of β-arr2 and Dvl2 expression was confirmed by immunoblot (Fig. 5A and B). APC stimulated a significant increase in SphK1 activity in endothelial cells transfected with non-specific siRNA (Fig. 5A), whereas aPC-induced SphK1 activity was markedly reduced upon β-arr2 depletion (Fig. 5A). In contrast, aPC-mediated activation of SphK1 remained unperturbed in Dvl2-depleted endothelial cells compared to non-specific siRNA transfected control cells (Fig. 5B), suggesting that β-arr2 functions as a key effector of aPC/PAR1-induced SphK1 activation. These findings further suggest that aPC stimulates divergent β-arr2-dependent cytoprotective signaling pathways.

**Figure 5.**
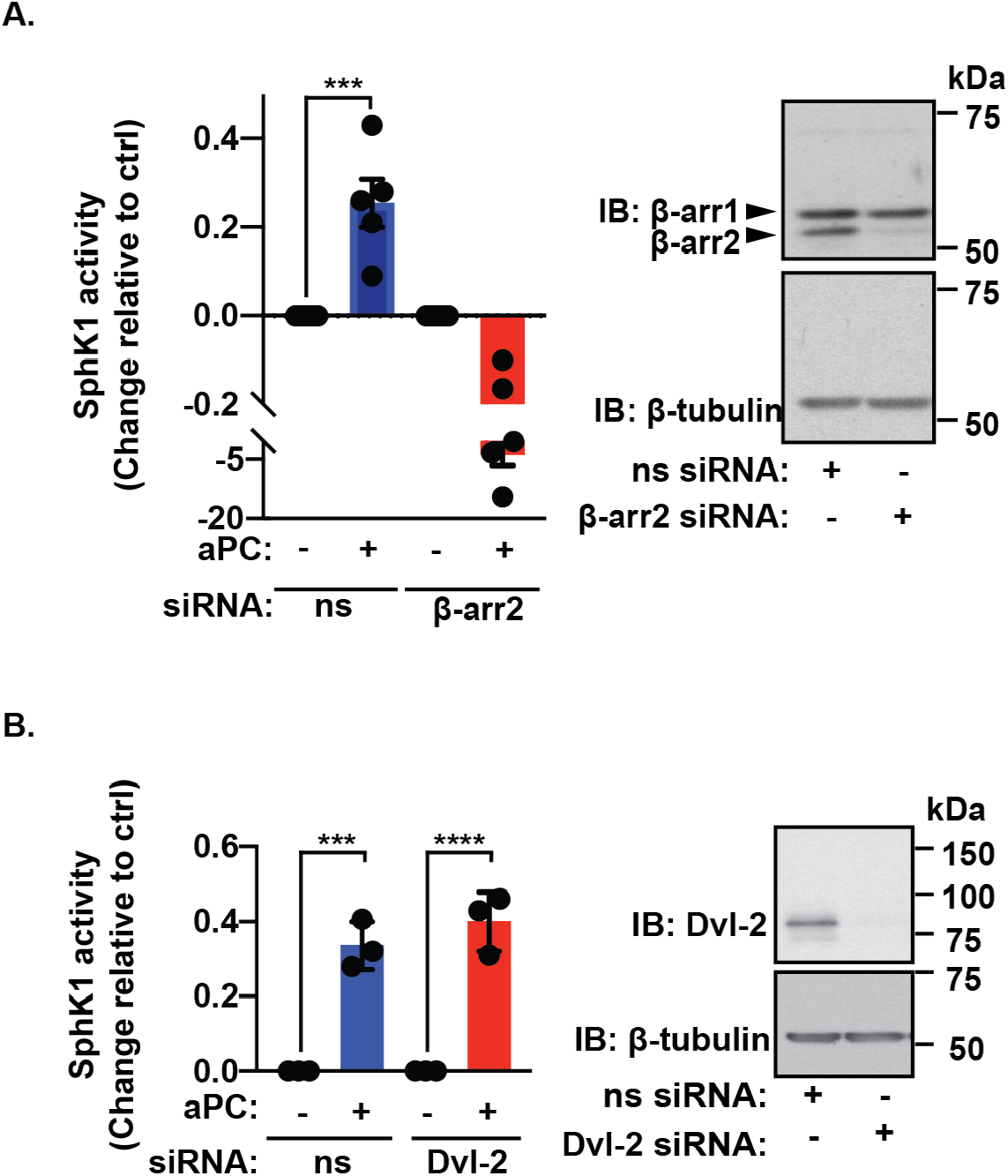
β-arr2 and not Dvl-2 mediates aPC-induced SphK1 activity. (A) EA.hy926 cells were treated with either non-specific (ns) siRNA or β-arr2 siRNA, serum-starved and then treated with 20 nM of aPC for 15 min. Cells were lysed and SphK1 activity was assessed. Data (mean ± S.D., n = 5) was analyzed by Student’s t-test. β-arr2 knockdown was verified by immunoblotting. (B) EA.hy926 cells were treated with either ns siRNA or Dvl-2 siRNA and SphK1 activity determined. Dvl-2 knockdown was confirmed by immunoblotting. Data (mean ± S.D., n = 5) was analyzed by Student’s t-test, ****, *P*<0.0001 and ***, *P*<0.001.

Next, we determined whether aPC/PAR1 transactivation of S1PR1-mediated Akt signaling is dependent on β-arr2 using siRNA-targeted depletion. Depletion of β-arr2 by siRNA was confirmed by immunoblotting (Fig. 6A and B, *insets*). APC stimulated a robust increase in Akt activation in non-specific siRNA transfected endothelial EA.hy926 cells (Fig. 6A, lanes 1-4), which was significantly inhibited in cells deficient in β-arr2 expression (Fig. 6A, lanes 5-8). The loss of β-arr2 failed to perturb aPC-induced ERK1/2 activation (Fig. 6B), indicating that aPC/PAR1-dependent β-arr2 function is specific to the Akt signaling pathway. In HUVECs, aPC-promoted Akt activation was also significantly inhibited in β-arr2 deficient cells compared to non-specific siRNA control cells (Fig. 6C, lanes 1-4 versus 5-8). In contrast, aPC-induced ERK1/2 activation was similarly robust in β-arr2 and non-specific siRNA transfected EA.hy926 cells and HUVECs (Fig. 6D, lanes 1-6 versus 7-12), suggesting that β-arr2 regulation of Akt is specific. To assess the function of β-arr2 in aPC-mediated pro-survival, endothelial cells were depleted of β-arr2 and aPC-mediated protection against TNF-α induced cell death was determined using Annexin V-FITC staining and flow cytometry. APC pretreatment significantly reduced TNF-α-induced cell death in non-specific siRNA transfected cells (Fig. 6E), whereas aPC protection against TNF-α-induced cell death was significantly diminished in β-arr2 deficient cells (Fig. 6E), consistent with a critical role for β-arr2 in aPC-mediated pro-survival. Taken together, our novel findings indicate that β-arr2 functions as a key regulator of aPC/PAR1-induced anti-apoptotic responses by driving a distinct SphK1-SPR1-Akt-mediated pro-survival signaling pathway (Fig. 7).

**Figure 6.**
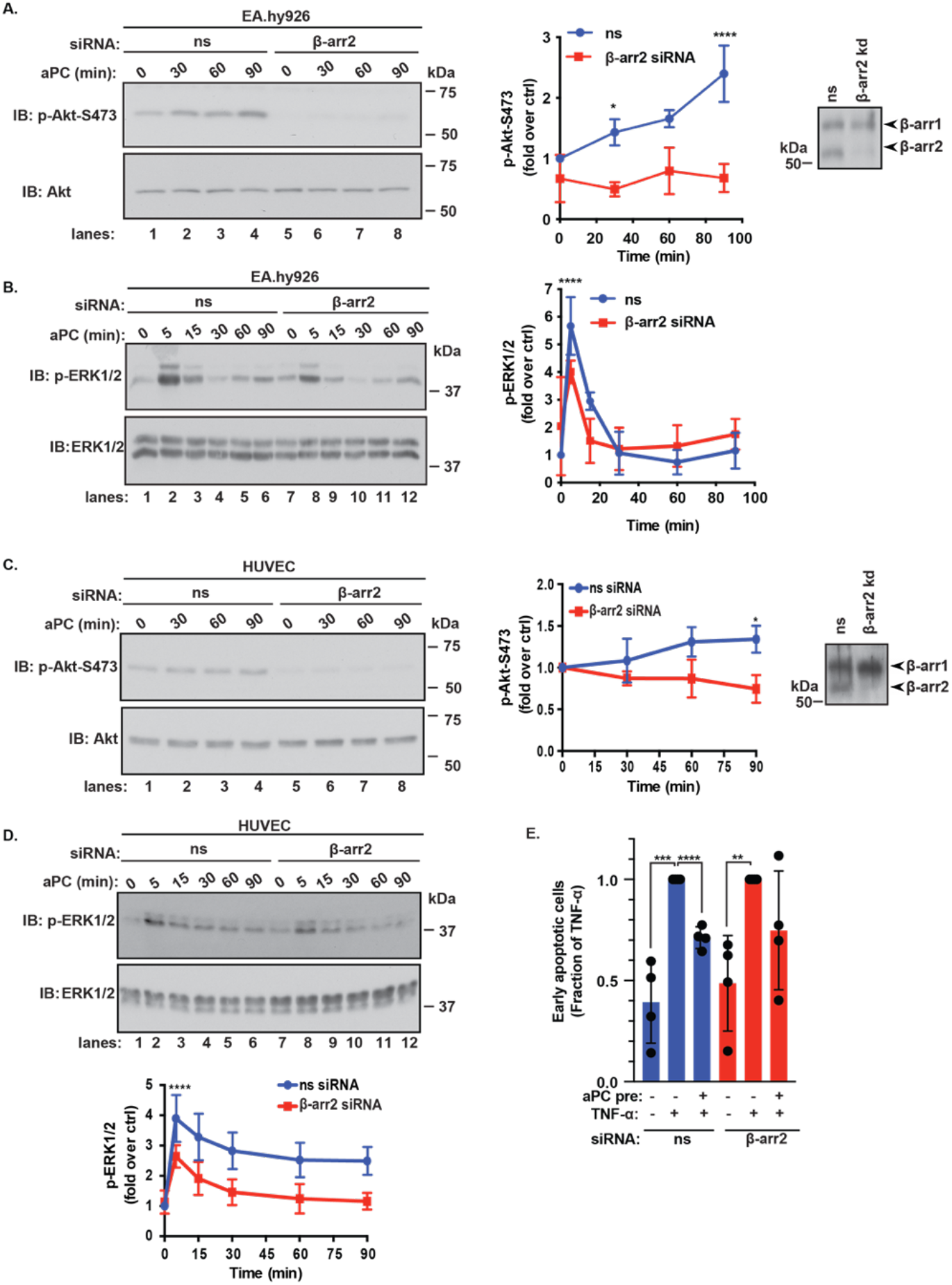
β-arr2 drives aPC anti-apoptotic responses. (A) EA.hy926 cells were transfected with non-specific (ns) or β-arr2 siRNA and stimulated with or without 20 nM aPC for various times, phospho-Akt-S473 was then detected by immunoblotting. Data (mean ± S.D., n = 3) was analyzed by two-way ANOVA. Cell lysates were immunoblotted to verify β-arr2 knockdown. (B) Cell lysates from A were immunoblotted for phospho-ERK1/2 and total ERK1/2. Data (mean ± S.D., n = 3) was analyzed by two-way ANOVA. (C) HUVECs were treated and phospho-Akt analyzed as described in A. (D) HUVECs were treated with aPC as described in A and ERK1/2 detected. (E) EA.hy926 cells transfected with ns or β-arr2 siRNA were stimulated with 20 nM aPC for 4 h and treated with 20 ng/mL TNF-α for 24 h and apoptotic response was determined. Data (mean ± S.D., n = 3) were analyzed by two-way ANOVA. ****, *P*<0.0001, ***, *P*<0.001, **, *P*<0.01 and *, *P*<0.05.

**Figure 7.**
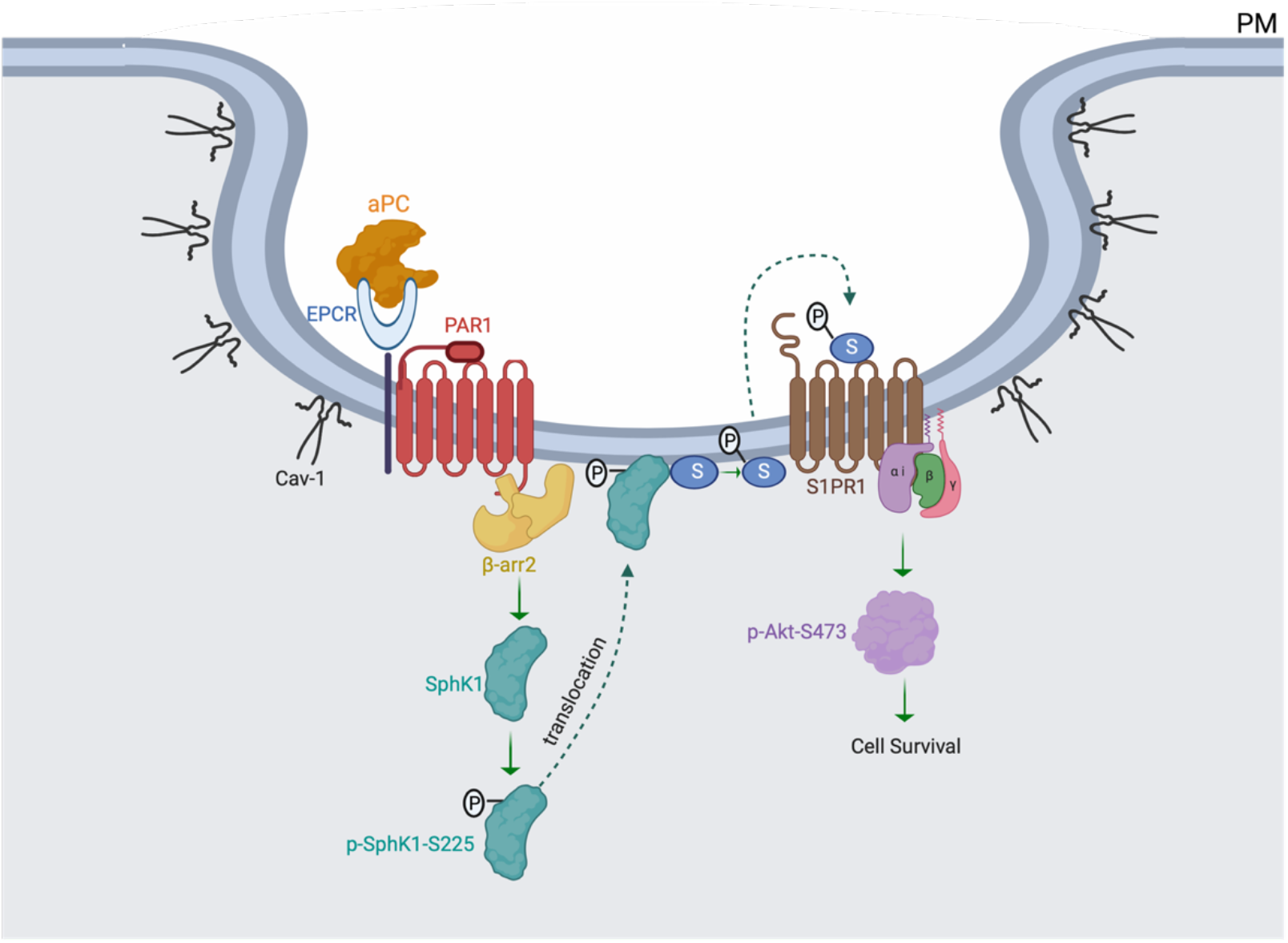
Model of aPC/PAR1 transactivation of S1PR1 via β-arr2-mediated SphK1 activation. Activation of PAR1 by aPC bound to its’ co-receptor EPCR promotes β-arr2-dependent SphK1 transactivation of S1PR1 that results in Akt at S473 phosphorylation, which protects endothelial cells from TNF-α induced cell death. SphK1 activation is associated with β-arr2-mediated phosphorylation, translocation to the plasma membrane following aPC stimulation.

## Discussion

Endothelial dysfunction results in barrier disruption and increased sensitivity to apoptosis. In this study we delineate a new GPCR-β-arr2 driven signaling pathway that regulates cellular resistance to apoptosis in the endothelium. We demonstrate that the aPC/PAR1-promoted anti-apoptotic response is mediated by Akt signaling induced via transactivation of the S1PR1 co-receptor. PAR1 and S1PR1 are required for aPC-induced protection against cell death, and both receptors reside in Cav-1 enriched microdomains. APC/PAR1-induced transactivation of S1PR1-Akt signaling is mediated by SphK1 activation. We further report that β-arr2 and not Dvl2 is required for aPC/PAR1-stimulated SphK1 activity. Moreover, β-arr2 functions as the key mediator of aPC/PAR1-induced endothelial cell survival by triggering the SphK1-S1PR1-Akt signaling pathway. Together, these studies reveal that aPC/PAR1-induced cytoprotective responses are mediated by discrete β-arrestin-2-driven signaling pathways that are modulated by co-receptors.

A major finding of our work is that aPC stimulates SphK1 activity via β-arr2 but not Dvl2, indicating that unique β-arr2-driven cytoprotective signaling pathways exist. Previous studies showed that aPC promotes barrier stabilization and protection against thrombin and vascular endothelial growth factor (VEGF)-induced barrier leakage (Claesson-Welsh et al., 2020; Rahimi, 2017). We discovered that aPC-activated PAR1 signals preferentially via β-arr2 and not heterotrimeric G proteins to confer protection against thrombin-induced barrier disruption (Soh & Trejo, 2011). We further demonstrated that aPC/PAR1 stimulates β-arr2-dependent polymerization of Dvl2 and Rac-1 activation, which facilitates endothelial barrier protection (Soh & Trejo, 2011). A recent study also showed that aPC occupancy of EPCR in endothelial cells promotes β-arr2- and Dvl2-mediated inhibition of cytokine-induced monocyte recruitment, an anti-inflammatory response (Roy et al., 2016). These studies suggest that β-arr2 and Dvl2 scaffolds might function as universal mediators of diverse aPC/PAR1 cytoprotective responses. However, we show that β-arr2 but not Dvl2 is essential for aPC-stimulated SphK1 activity, which is required for transactivation of S1PR1-induced Akt activity and anti-apoptotic responses. APC cytoprotective responses are well documented in other cell types such as neurons, podocytes or immune cells (Guo et al., 2004; Guo et al., 2013; Madhusudhan et al., 2012), but whether these responses are also uniquely regulated by distinct β-arr2-mediated signaling pathways is not known.

Caveolae organize cellular signal transduction by bringing effectors in close proximity to receptors through binding to Cav-1 (Fridolfsson et al., 2014). A unique feature of aPC/PAR1 cytoprotection is the localization of key signaling components, including PAR1, EPCR, and β-arr2 in caveolae (Bae et al., 2007, 2008; Soh & Trejo, 2011). Previous studies have shown that caveolae are required for aPC/EPCR/PAR1 complex formation and β-arr2-mediated cytoprotection (Bae et al., 2007, 2008; Rezaie, 2011; Russo et al., 2009; Soh & Trejo, 2011). Here, we now report that S1PR1 also resides in caveolae together with PAR1. In addition, aPC stimulation of endothelial cells caused a significant increase in S1PR1 binding to Cav-1 via a PAR1-dependent mechanism. Although, it is unclear whether S1PR1 binds directly to Cav-1 or other components such as EPCR or PAR1 after recruitment to caveolae. Previous studies showed that oxidized lipids promote Cav-1 phosphorylation and enhance recruitment of S1PR1, Akt, and other effectors to caveolae in endothelial cells (Karki et al., 2018; Singleton et al., 2009). Another report demonstrated that aPC stimulates S1PR1 phosphorylation (Chavez et al., 2015), but neither study assessed the function of phosphorylation on EPCR and S1PR1 co-association or interaction with Cav-1. Our studies demonstrate that PAR1 and S1PR1 co-exist in caveolae and co-localize at the cell periphery; however, aPC-stimulation did not induce internalization of either receptor. While our work demonstrates that S1PR1 remains at the cell surface following aPC stimulation, it is unclear whether the phosphorylation state of S1PR1 is changed. Thus, additional studies are needed to understand how Cav-1 regulates aPC-dependent cytoprotective signaling, particularly of the mechanisms that mediate S1PR1 interaction with caveolin-1 and/or recruitment to caveolae.

Our study further highlights diverse functions of co-receptors in mediating aPC/PAR1 cytoprotective signaling. S1PR1 is not the only co-receptor that is required for aPC/PAR1-induced cytoprotective activities. aPC/PAR1 activation of Akt requires distinct co-receptors that are cell type-dependent (Feistritzer & Riewald, 2005; Finigan et al., 2005; Guo et al., 2013). In endothelial cells, aPC-stimulated Akt phosphorylation and anti-apoptotic responses are also dependent on Apolipoprotein E Receptor 2 (ApoER2) (Sinha et al., 2016), but a link between Akt-mediated apoptosis was not established. This study further showed that ApoER2 mediates aPC-induced Rac-1 activation and barrier stabilization (Sinha et al., 2016), suggesting that multiple co-receptors mediate aPC cytoprotective responses. In neurons, aPC/PAR1 promotes Akt-mediated neuronal proliferation, migration, differentiation and apoptosis via S1PR1 and PAR3 co-receptors (Guo et al., 2013). Previous studies showed that aPC cleaves and activates PAR3 and aPC-derived PAR3 tether-ligand peptides protects against VEGF-induced endothelial barrier hyperpermeability (Burnier & Mosnier, 2013; Heuberger et al., 2019) In kidney podocytes, aPC inhibits apoptosis through a PAR3-dependent mechanism (Madhusudhan et al., 2012). In addition, PAR3 dimerization with PAR2 in human podocytes and with PAR1 in mouse podocytes facilitates aPC-mediated protection against apoptosis. PAR3 dimerization with PAR1 and PAR2 is further regulated by Cav-1 association (Madhusudhan et al., 2012). The Tie2 receptor tyrosine kinase also interacts with aPC to promote endothelial barrier stabilization (Minhas et al., 2010; Minhas et al., 2017). However, the impact of co-receptors on aPC/PAR1-induced downstream signaling effectors such as β-arr2 and Dvl2 in distinct cell types has not been thoroughly examined.

GPCR signaling is diverse, complex, and occurs in various cellular compartments on the plasma membrane and on endosomes to orchestrate an effective inflammatory response (Birch et al., 2021). Similarly, aPC/EPCR/PAR1 cytoprotective signaling is diversely regulated by different co-receptors in distinct cell types to yield critical cytoprotective functions. However, the molecular effectors important for conveying aPC/PAR1 downstream cytoprotective signaling in most cell types are largely unknown. We conducted a global phosphoproteomic analysis of aPC-stimulated endothelial cells to identify pathways and proteins that confer aPC/PAR1 biased signaling. This work led to the identification of Adducin-1 as a key regulator of aPC-stimulated Akt signaling (Lin et al., 2020). The aPC phosphoproteome also revealed significant enrichment of proteins associated with the nucleus, mRNA splicing, DNA binding and chromatin regulators. Actin binding was also enriched as well as phosphopeptides associated with adherens junctions was further evident, likely related to aPC’s role in enhancing endothelial barrier stabilization (Lin et al., 2020; Russo et al., 2009; Soh & Trejo, 2011). Thus, a rich array of proteins associated with biological processes induced by aPC exist and are available for further interrogation. In summary, this study demonstrates that distinct aPC/PAR1 cytoprotective responses are driven by discrete β-arr2-mediating signaling pathways that are specifically modulated by different co-receptors in endothelial cells.

## Materials and methods

### Cell culture

EA.hy926 cells (ATCC, #CRL-2922) were grown at 37°C, 8% C0_2_ in 10% FBS-DMEM (Gibco, #10-013-CV and #10437-028) supplemented with fresh 20% pre-conditioned media every two days and cultured to passage 8. Pooled primary HUVECs (Lonza, #C2519A) were grown at 37°C, 5% C0_2_ in EGM-2 (Lonza, #CC-3162), media was changed every two days and cultured to passage 6. EA.hy926 and HUVECs were grown for 4 to 5 days until confluence and then incubated overnight in 0.4% FBS-DMEM. Cells were then washed and serum-starved in DMEM containing 10 mM HEPES, 1 mM CaCl_2_, and 1 mg/mL BSA for 1 h prior to agonist, antagonist, and inhibitor treatments as described below.

### Inhibitor and antagonist treatments

Serum-starved cells were preincubated at 37°C with 100 nM PF-543 (Tocris, #5754) or 10 µM W146 (Tocris, #3602) for 30 min, or with 1 µM MK-2206 (Selleck, #S1078) or 10 µM Vorapaxar (Axon Medchem, #1755) for 1 h.

### Transfections with siRNAs

Cells were seeded at 1.4×10^5^ cells per well in a 12-well plate and grown as described above. Cells were transfected with siRNA using the TransIT-X2 System (Mirus, #MIR 600) according to the manufacturer’s instructions. The following siRNAs were used: 50 nM β-arrestin-2 siRNA (Dharmacon) 5’-GGACCGCAAAGTGTTTGTG-3’, 12.5 nM Dvl2 siRNA #2 (Qiagen, #SI00063441) 5’-CACGCTAAACATGGAGAAGTA-3’, 25 nM of S1PR1 #1 (Qiagen, #SI00376201) 5’-ATGATCGATCATCTATAGCAA-3’ and 25nM S1PR1 #2 (Qiagen, #SI00376208) 5’-TAGCATTGTCAAGCTCCTAAA-3’ siRNAs or AllStars Negative Control siRNA (Qiagen, #1027281). After 72 h, protein expression was determined by immunoblotting using anti-β-arr2 A2CT (a generous gift from Dr. Robert Lefkowitz, Duke University), anti-Dvl2 (CST, #3216), anti-S1PR1 (Santa Cruz Biotechnology, #sc-25489), anti-GAPDH (GeneTex, #GTX627408), and anti-tubulin (CST, #86298S) antibodies.

### Cell death assays and flow cytometry

EA.hy926 cells were plated at 1.4×10^5^ cells per well in a 12-well plate. Serum-starved cells were pretreated with 20 nM aPC (Hematologic Technologies, #HCAPC-0080) for 4 h and then incubated with 25 ng/mL TNF-α (PeproTech, #300-01A) for 24 h, or post-treated with 20 nM aPC for either 1, 2, or 3 h. Cells were gently harvested using Cellstripper (Corning, #25-056-CL) and washed with cold PBS. Cells were resuspended in 40 μL 1X Annexin V binding buffer (Biolegend, #422201) + 2 μL Annexin V FITC (BioLegend, #640906) and incubated at room temperature for 15 min, protected from light. Cells were washed with 160 μL of 1x Annexin V binding buffer, followed by a 5 min centrifugation at 550g. Pelleted cells were resuspended in 200 μL 1X Annexin V binding buffer + 2 uL of 100 μg/mL propidium iodide (PI) (Sigma-Aldrich, #P4170). Data acquisition was performed on a BD FACS Canto II Flow cytometer (BD Biosciences) on a log scale with 30,000 singlet gate events collected per sample. Data compensation and analysis were performed with FlowJo v10 software (Tree Star). The gating strategy was as follows: Annexin V and PI negative events were backgated to FSC-A/SSC-A to determine cell debris. A “not gate” was made based on cell debris in FSC-A/SSC-A. Doublet discrimination was performed using FSC-A vs FSC-H and SSC-A vs SSC-H. The resulting gated cells were analyzed for Annexin and PI staining and reported as percent of singlets.

### Assay for detecting TNF-α-induced caspase-3 cleavage

EA.hy926 cells were seeded at 6.2×10^5^ cells per well in a 6-well plate. Serum-starved cells were pretreated with or without 20 nM aPC for 3 h followed by treatment with 25 ng/mL TNF-α for 24 h. Cells were washed with PBS and lysed in RIPA buffer (50 mM Tris HCl, pH 8.0,150 nM NaCl, 1% NP-40, 0.5% Sodium Deoxycholate, 0.1% SDS) supplemented with protease inhibitors (1 mM PMSF, 2 µg/mL Aprotinin, 10 µg/mL Leupeptin, 1µg/mL Pepstatin, and 1 µg/mL Trypsin protease inhibitor). Cells were sonicated at 10% amplitude for 10 sec and clarified by centrifugation at 20,817g for 15 min. Cell lysates were subject to immunoblotting with anti-Caspase-3 (CST, #9662S), anti-Cleaved-Caspase-3 (CST, #9661), and anti-GAPDH antibodies, followed by secondary anti-mouse or anti-rabbit HRP conjugated antibodies (Bio-Rad, #170-6516 and #170-6515) and quantified by densitometry analysis using ImageJ software.

### Signaling assays

EA.hy926 cells were seeded in a 24-well plate at 1.4×10^5^ cells per well. HUVECs were seeded at 1.77×10^5^ cells per well in a 24-well plate. Serum-starved cells were treated with either 20 nM of aPC or 1 µM of S1P (Tocris, #1370) for various times, and then lysed in 2X Laemmli Sample Buffer (LSB) containing 200 mM Dithiothreitol (DTT). Equivalent amounts of cell lysates were immunoblotted with anti-SphK1-S225 (ECM Biosciences, #SP1641), anti-SphK1 (ECM Biosciences, #SP1621), anti-p-Akt-S473 (CST, #4060), anti-Akt (CST, #9272S), anti-p-ERK1/2 (CST, #9106L), anti-ERK1/2 (CST, #9102L) antibodies followed by secondary anti-mouse or anti-rabbit HRP conjugated antibodies. Immunoblots were quantified as described above.

### Membrane and cytosolic subcellular fractionation

EA.hy926 cells were seeded at 2.7×10^6^ cells per 15 cm dish. Serum-starved cells were treated with or without 20 nM aPC for 1 h at 37°C. Cells were washed 2X with cold PBS, lysed in fractionation buffer (250 mM Sucrose, 20 mM HEPES, 10 mM KCl, 1.5 mM MgCl_2_, 1 mM EDTA, 1 mM EGTA, 1 mM DTT, β-glycerophosphate, 50 mM NaF, and 1 mM Okadaic acid) and lysates were gently passed through a 25G needle. Cell lysates were subjected to several sequential centrifugations at 4°C. Lysates were centrifuged at 106g for 3 min and the pellet used as the nuclear sample. Supernatant was centrifuged at 6797g for 10 min, and pellet was saved as the mitochondrial fraction. Supernatant was ultracentrifuged at 100,000g for 1 h, and the supernatant used as the cytosolic fraction and the pellet membrane fraction. The membrane pellet was resuspended and homogenized by passing through a 25G needle. The membrane pellet was then ultracentrifuged at 100,000g for 45 min. All pellets were washed, resuspended, and sonicated. Lysates were resuspended in LSB, boiled, and immunoblotted with anti-SphK1 (ECM Biosciences, #SP1621), anti-GAPDH (GeneTex, #GTX627408), and Na^+^/K^+^ ATPase (CST, #3010) antibodies.

### SphK1 activity assay

EA.hy926 cells were seeded in 6-well plates at 6×10^5^ cells per well. Serum-starved cells stimulated with 20nM aPC for 15 min, washed with cold PBS and SphK1 activity assay performed according to the manufacturer instructions (Echelon, #K-3500). In brief, cells were resuspended in Reaction Buffer with 1 mM of DTT and sonicated for 10 sec at 10% amplitude. Total protein in cell lysates was quantified using the bicinchoninic acid (BCA) protein assay (ThermoFisher, #23221 and #23224) and normalized to 1.5 mg/ml of protein for each sample. Then, 400 mM of sphingosine solution and 10 µL of each sample were aliquoted into each well of a 96-well plate. The reaction was initiated with the addition of 20 µM of ATP to each sample, incubated for 30 min, followed by the addition of K-LUMa ATP detector per well for 10 min to stop the reaction. Luminescence was determined using the Tristar LB 941 Plate Reader (Berthold Technologies). A reduction in luminescence compared to control indicates ATP depletion or consumption and used as an assessment of SphK1 Activity. To generate positive or negative values for increased or reduced SphK1 activity respectively, background luminescence was subtracted from the raw luminescence units (RLUs) for each sample. Using an ATP standard curve, the concentration of ATP after the 30 min reaction was determined then subtracted from the starting ATP concentration of 20 µM. This yielded the concentration of ATP consumed in 30 min. The difference from control values were plotted for each sample with three or more replicates for each experiment.

### Immunofluorescence confocal microscopy

EA.hy926 cells were plated on coverslips in 12-well plate at a density of 1.4×10^5^ cells per well. Serum-starved cells were stimulated with or without 20 nM of aPC for 1 h or 10 nM ±-thrombin for 1 h (Enzyme Research Laboratories, #HT 1002a), washed with cold PBS, and incubated with PBS for 10 min. Endogenous PAR1 was labeled with anti-PAR1 WEDE antibody (Beckman Coulter, #IM2584) at 1:500 for 1 h on ice, cells were treated with or without agonist, fixed for 5 min with 4% paraformaldehyde and permeabilized with 0.1% Triton-X 100. The detection of S1PR1 was determined using anti-S1PR1 antibody (Santa Cruz Biotechnology, #sc-25489) diluted at 1:100 in 0.03% BSA, 0.01% Triton-X 100 and 0.01% normal goat serum overnight at 4°C. Secondary fluorescent antibodies anti-mouse-Alexa-488 (Invitrogen, #A-11001 and anti-rabbit-Alexa-594 (Invitrogen, #A-11012) were diluted at 1:750 incubated at room temperature for 1 h in 0.03% BSA, 0.01% Triton-X 100 and 0.01% normal goat serum. Slides were mounted using ProLong Gold Antifade Mountant (Invitrogen, #P10144). Confocal images were acquired using sequentially using the same settings with an Olympus IX81 spinning-disk Microscope (Tokyo, Japan) equipped with a CoolSNAP HQ2 CCD Camera (Andor) and 63x Plan Apo objective (1.4 NA) with appropriate excitation-emission filters. Line-scan analysis was performed using Image J software (NIH, Maryland, USA).

### Immunoprecipitation assays

EA.hy926 wildtype cells and EA.hy926 cells stably expressing PAR1-specific shRNA pSilencer Retro (Russo et al., 2009) were grown in 10 cm dishes, serum-starved overnight, treated with 20 nM aPC, and lysed in Triton-X 100 lysis buffer (50 mM Tris-HCl, pH 7.4, 100 mM NaCl, 10 mM NaF, 1% Triton-X 100 supplemented with protease inhibitors). Cell lysates were homogenized, clarified by centrifugation and protein concentrations determined by BCA. Equivalent amounts of lysates were subjected to immunoprecipitations using the anti-S1PR1 (Santa Cruz Biotechnology, #sc-25489) and anti-PAR1 WEDE antibodies. Immunoprecipitates were resuspended in 2X LSB containing 200 mM DTT, and eluents immunoblotted using S1PR1 (Santa Cruz Biotechnology, # sc-25489), Cav-1 (CST, #610060), and PAR1 antibodies (Beckman Coulter, #IM2584) and developed by chemiluminescence.

### Sucrose fractionation

EA.hy926 cells were plated at 4.95×10^6^ cells per 10 cm dish. Cells were washed with cold PBS, lysed in sodium carbonate buffer (150 mM sodium carbonate, pH 11, 1 mM EDTA, supplemented with protease inhibitors) with a dounce homogenizer, passed through 18G needle 10X, and sonicated on ice at 10% amplitude. Cell lysates were mixed with equal volume of 80% sucrose in MES-buffered saline (25 mM MES pH 6.5, 150 NaCl, and 2 mM EDTA) supplemented with 300 mM sodium carbonate for a total of 1.6 mL in a 12 mL ultracentrifuge tube (Beckman Coulter, #343778). Approximately, 6 mL of 35% MES-buffered saline supplemented with 150 mM sodium carbonate was added to the top of the tube gently without perturbing solution on the bottom, and 4 mL of 5% sucrose in MES-buffered saline supplemented with 150 mM sodium carbonate on top of the 35% MES-buffered saline solution. Samples were placed in SW41 rotor and ultracentrifuged for 18-20 h at 4°C, at 229884g. The 1 mL fractions were collected sequentially, and samples were immunoblotted using anti-S1PR1 (Santa Cruz Biotechnology, # sc-25489), EEA1 (BD Biosciences, #610457), anti-PAR1 WEDE (Beckman Coulter, #IM2584), and anti-Cav-1 (CST) antibodies.

## Data analysis

Data were analyzed with Prism 9.0 statistical software. Statistical analysis methods are indicated in the Figure legends.

## Author contributions

Olivia Molinar-Inglis and Cierra Birch, Conceptualization, Investigation, Writing—original draft, Writing—review, and editing; Dequina Nicholas, Metztli Cisneros-Aguirre, Anand Patwardhan, Buxin Chen, Neil J. Grimsey, Patrick G. Menzies, Huilan Lin, Luisa J. Coronel, Mark A. Lawson, Hemal. H. Patel, Investigation; JoAnn Trejo, Conceptualization, Investigation, Writing—review and editing.

## Acknowledgements

We thank members of the Trejo lab for advice and guidance. We also thank Dr. Antonio De Maio (UC San Diego) for assistance with Flow Cytometry and Cara R. Rada and Hilda Mejia-Peña assisting with HUVEC cell culture. This work was supported by NIH/NIGMS R01 GM116597 and R35 GM127121 (J.T.), NIH/NIGMS R25 GM083275 (M.C., L.J.C.), NIH/NHLBI T32 HL007444 (C.B.), NIH/NIGMS K12 GM068524 (O.M.I., D.N.), NIH/NICHD P50 HD12303 (M.A.L), and UC President’s Postdoctoral Fellowship (O.M.I. and D.N.).

## Competing interest statement

The authors declare no competing interests

---

**Supplemental Figure 1.**
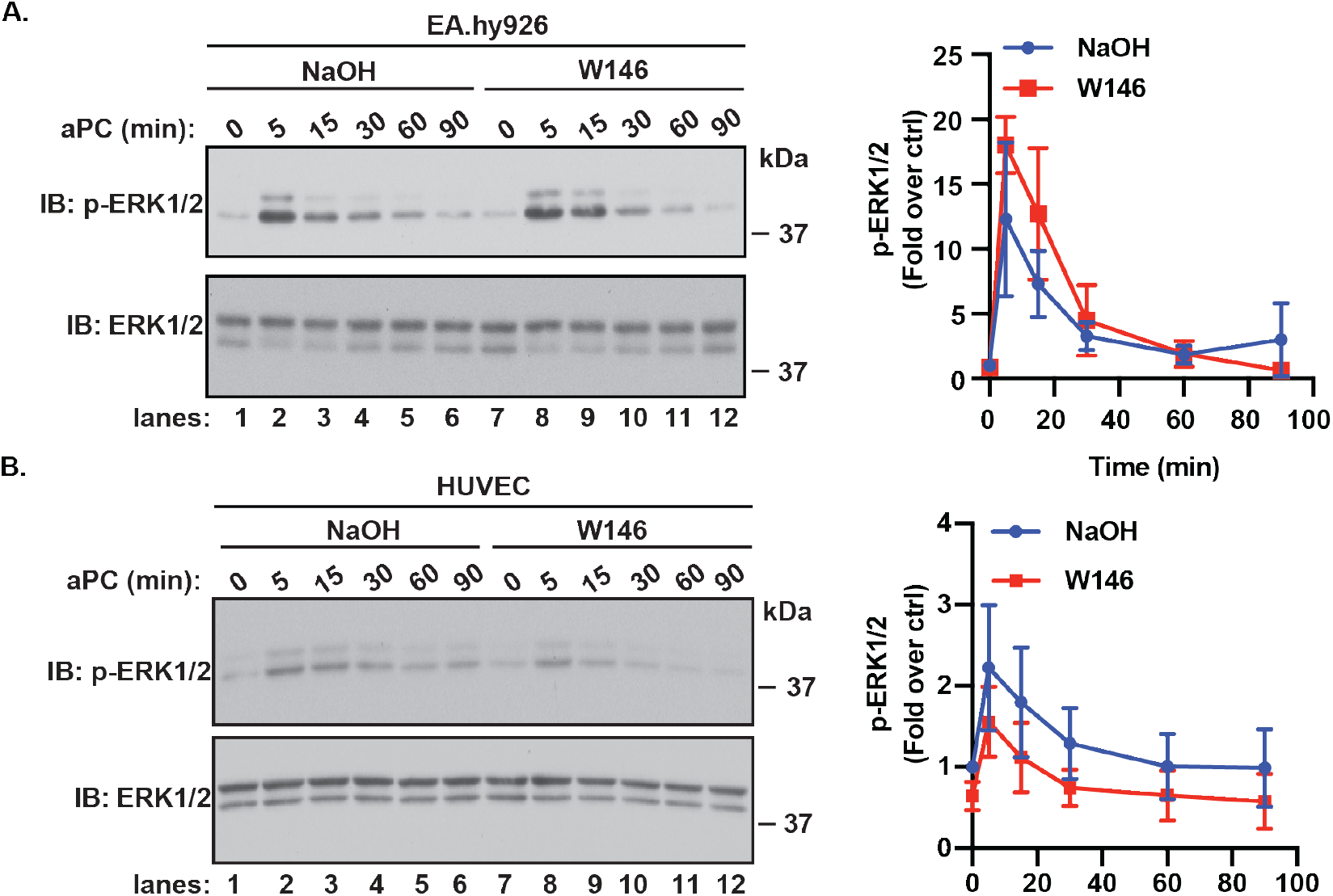
S1PR1 is not involved in modulating the aPC-PAR1-ERK1/2 axis. A) EA.hy926 cells were pretreated with or without 10 µM W146 and stimulated with or without 20 nM of aPC for various times. Phospho-ERK1/2 was detected by immunoblotting. Data (mean ± S.D., n = 3) was analyzed by two-way ANOVA. (B) Same as A but in HUVECs.

**Supplemental Figure 2.**
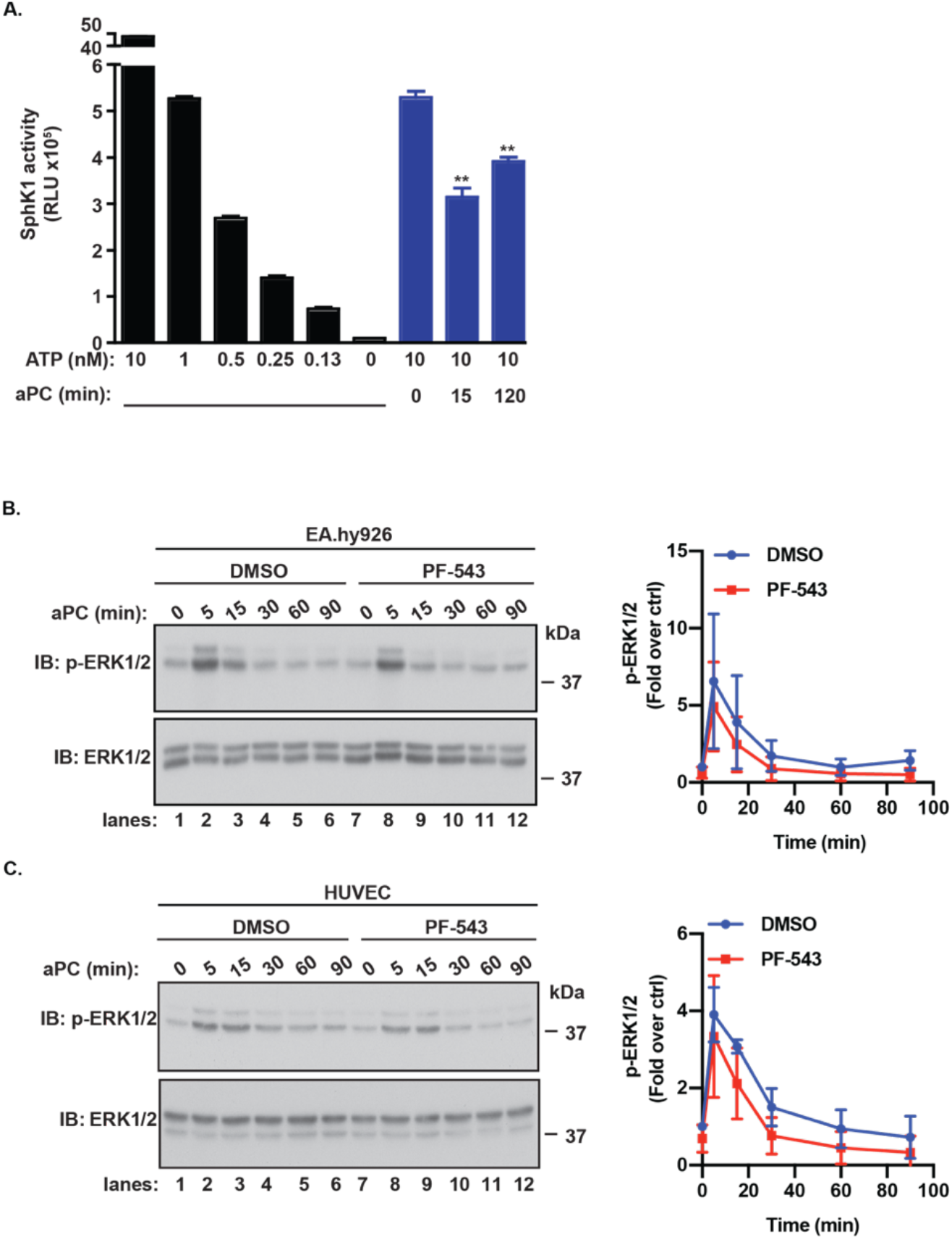
ATP standard curve for SphK1 activity. (A) EA.hy926 cells were treated with 20 nM aPC for various times. Cells were lysed and SphK1 activity was measured using an Echelon SphK1 activity assay. The luminescence of an ATP standard curve and aPC treated cells was deteremined and the raw values are shown. Data (mean ± S.D., n = 3) was analyzed by Student’s t-test. **, *P*<0.001. (B) EA.hy926 cells were pretreated with or without 100 nM PF-543 and stimulated with or without 20 nM of aPC for various times. Phospho-ERK1/2 was detected by immunoblotting. Data (mean ± S.D., n = 3) was analyzed by two-way ANOVA. (B) Same as A but in HUVECs.

## Notes

### Competing Interest Statement

The authors have declared no competing interest.

### Summary of Updates

Figure 3 revised, mistakenly duplicated Figure 4

